# Neurogranin Stimulates Ca^2+^/calmodulin-dependent Kinase II by Inhibiting Calcineurin at Specific Calcium Spike Frequencies

**DOI:** 10.1101/597278

**Authors:** Lu Li, Massimo Lai, Stephen Cole, Nicolas Le Novère, Stuart J. Edelstein

## Abstract

Calmodulin sits at the centre of molecular mechanisms underlying learning and memory. Its complex, and sometimes opposite influences, via the binding to various proteins, are yet to be fully understood. Calcium/calmodulin-dependent protein kinase II (CaMKII) and calcineurin (CaN) both bind open calmodulin, favouring Long-term potentiation (LTP) or depression (LTD) respectively. Neurogranin binds to the closed conformation of calmodulin and its impact on synaptic plasticity is less clear. We set up a mechanistic computational model based on allosteric principles to simulate calmodulin state transitions and its interaction with calcium ions and the three binding partners mentioned above. We simulated calcium spikes at various frequencies and show that neurogranin regulates synaptic plasticity along three modalities. At low spike frequencies, neurogranin inhibits the onset of LTD by limiting CaN activation. At intermediate frequencies, neurogranin limits LTP by precluding binding of CaMKII with calmodulin. Finally, at high spike frequencies, neurogranin promotes LTP by enhancing CaMKII autophosphorylation. While neurogranin might act as a calmodulin buffer, it does not significantly preclude the calmodulin opening by calcium. On the contrary, neurogranin synchronizes the opening of calmodulin’s two lobes and promotes their activation at specific frequencies, increasing the chance of CaMKII trans-autophosphorylation. Taken together, our study reveals dynamic regulatory roles played by neurogranin on synaptic plasticity, which provide mechanistic explanations to opposing experimental findings.

**Author Summary:** How our brains learn and remember things lies in the subtle changes of the strength with which brain cells connect to each other, the so-called synaptic plasticity. At the level of the recipient neuron, some of the information is encoded into patterns of intracellular calcium spikes. Calmodulin, a small bilobed protein which conformation is regulated by the binding of calcium ions, decodes these signals, and modulates the activity of specific binding partners.

Two key regulators, calcineurin and calcium/calmodulin-dependent protein kinase II, which respectively weaken or strengthen synaptic connections, bind both lobes of calmodulin in its open form, favoured by calcium. On the contrary, neurogranin binds preferentially to one lobe of calmodulin, in the closed form, unfavoured by calcium. It was thus initially suggested that it would inhibit calmodulin activation and decrease synaptic plasticity. However, past research showed that neurogranin sometimes actually enhances synaptic plasticity, though the mechanism was unclear.

Our computational models showed that neurogranin synchronizes the activation of the two lobes of calmodulin, favouring opening at high frequency calcium spikes. By doing so, neurogranin increases the impact of calmodulin on calcium/calmodulin-dependent protein kinase II and reduces its effect on calcineurin, resulting in a strengthening of synaptic connections.

## Introduction

Calmodulin is a small ubiquitous calcium-binding protein which cellular activity is mediated via the binding of a large array of target proteins. Calmodulin comprises two lobes, each binding two calcium ions, that can undergo transitions between a closed (T) and an open (R) conformations. Calcium binding shifts the transition towards the open form [1, 2]. Some calmodulin-binding proteins favour the open conformation while others tend to bind preferentially to the closed conformation [3–8].

In neurons, calmodulin plays a crucial role in mediating calcium regulation of synaptic plasticity, a mechanism underlying learning and memory. Among its main binding partners are Ca^2+^/calmodulin-dependent protein kinase II (CaMKII), and calcineurin (CaN), which both preferably bind to the open (R) form of calmodulin, and compete to facilitate long-term potentiation(LTP) or long-term depression (LTD) respectively [2, 9, 10]. On the contrary, neurogranin (Ng), an IQ domain-containing protein, binds preferentially to the closed (T) state, therefore, sequestering calmodulin from its targets in the absence of calcium [11–13]. Ng’s role in the regulation of synaptic plasticity is still under debate. Ng seems to facilitate LTP induction at high frequency electrical pulses [14–18]. However, it has also been reported to produce the opposite effect [19]. Deciphering the mechanisms underlying Ng’s opposing functions, and importantly, how Ng, CaMKII and CaN coordinate their activities in response to various signals, is crucial for understanding how calmodulin regulates brain function and dysfunction.

Although structurally similar, the two calmodulin lobes display different properties. The carboxy-terminal (C) lobe possesses higher calcium binding affinities but slow kinetics while the amino-terminal (N) lobe has lower calcium affinities but faster binding kinetics [6, 20–23]. In addition, the lobes exhibit a significant degree of structural autonomy [24, 25]. Calmodulin lobes undergo constant transitions between the open and closed conformations, transitions regulated by binding of both calcium ions and target proteins. The key to understand calmodulin function lies in the mechanisms underlying the differential activation of its targets and how binding these targets feedbacks to its conformational changes.

The two lobes contribute unevenly to targets binding. Targets binding preferentially to the closed state, especially Ng, interact predominantly with the C lobe [5, 7, 13, 26–28], whereas targets binding preferentially to the open state, such as CaMKII and CaN, contact both domains [4, 6, 8, 29, 30]. Accordingly, binding these targets affect differentially the conformational shifts of both lobes, as indicated by the modification of apparent calcium binding affinities [5, 6, 21, 26], targets stabilising the conformation for which they have the highest affinity (i.e. the complexes have the lowest free energy). Reciprocal influences between target binding and calcium binding often exhibit lobe specificity as well [5, 31–33]. Therefore, how calmodulin regulates the function of its binding partners, including CaMKII, CaN and Ng, is tightly linked with its structure and the functional differences between lobes.

Number of computational models have been developed to study the regulation of bidirectional synaptic plasticity [34–39], in which calmodulin activation is modelled either phenomenologically, using Hill or Adair-Klotz equations, or using sequential binding of calcium ions. These model disregarded calcium binding site specificities or lobe differences because their focus was mostly on the activation/autophosphorylation of CaMKII as the main regulator of synaptic plasticity. We proposed models based on a mechanistic description of calmodulin within an allosteric framework [2, 10], but that did not take the role of Neurogranin into account. A few models attempted to explain how Ng facilitates LTP, putting forward the buffering of calmodulin by Ng in the post-synaptic density (PSD) and the reshaping of free calcium spikes [40, 41]. Using mathematical modelling, Romano et al [42] recently showed that Ng facilitates CaMKII activation at a 100 Hz-tetanus stimulation. However they failed to show why, at the same time, Ng hinders LTP induction, observed by the same experimental group [19]. All these models lacked detailed mechanistic descriptions of calmodulin function.

In this paper, we present a detailed mechanistic model of calmodulin in the context of synaptic plasticity and its interaction with CaMKII, CaN and Ng, based on the allosteric framework and our previous hemiconcerted calmodulin model [25]. In our model, the four calcium binding sites were explicitly modelled and each lobe of calmodulin undergoes independent state transitions. In addition, we systematically estimated the allosteric parameters concerning the two calmodulin lobes together, as well as the kinetic constants, based on steady states and calcium chelating time course experimental data. We simulated a wide range of calcium spike frequencies in a wild type context and for a neurogranin knock-out situation. We showed the differences of calmodulin lobe responses and how they contribute to, but also are influenced by, the differential activations of calmodulin’s binding targets. We show that Ng is not merely a calmodulin buffer protein, but it can adjust the activation of calmodulin at frequencies maximizing CaMKII activation. Ng regulation of LTP induction depends on the local concentration of CaN, and on the amount of CaMKII required for inducing LTP.

## Materials and Methods

### Model Structure

We built a mechanistic mathematical model to describe the conformational changes of calmodulin lobes, calcium bindings and calmodulin interaction with binding partners. This model is based on our previous hemiconcerted allosteric model of calmodulin by Lai *et al.* [25]. The major differences between our model and other published calmodulin models are: 1) We do not assume sequential bindings of calcium ions to the four binding sites. Instead, each calcium binding site on calmodulin is independent, and independent from the binding of protein partners. 2) Calcium saturation does not directly affect calmodulin’s affinity for its protein binding partners. Binding CaMKII, CaN or Ng does not necessarily results in increased calcium binding affinities. Instead, modifications of the affinity for all binding partners are solely derived from calmodulin conformational changes. Calcium saturation decreases the free energy of the open state, favouring the association with proteins preferentially binding to that conformation and hindering the association with proteins preferentially binding to the closed state. Symmetrically, protein binding partners shift the conformation equilibrium of calmodulin towards their preferred state. On top of the model developed previously by Lai *et al.* [25], We explicitly modelled calmodulin’s interaction with CaMKII, CaN and Ng. We detailed the autophosphorylation of CaMKII monomers. We modelled their dephosphorylation as if directly mediated by CaN, omitting the intermediate reactions involving DARPP-32/Inhibitor-1 and Protein Phosphatase 1. We also implemented reactions to maintain basal calcium concentration, as well as enabling calcium spikes at different frequencies, amplitudes and durations. Reactions were primarily encoded in mass-action law kinetics except for the reactions depicting the calcium pump and CaMKII dephosphorylation. The model structure is partially illustrated in Fig. 1, using no calcium or one calcium ion bound calmodulin as examples. The full model is written by the Systems Biology Markup Language (SBML) [43] in the supplementary material and has been deposited in BioModels [44] (accession number: MODEL1903010001).

**Fig 1.**
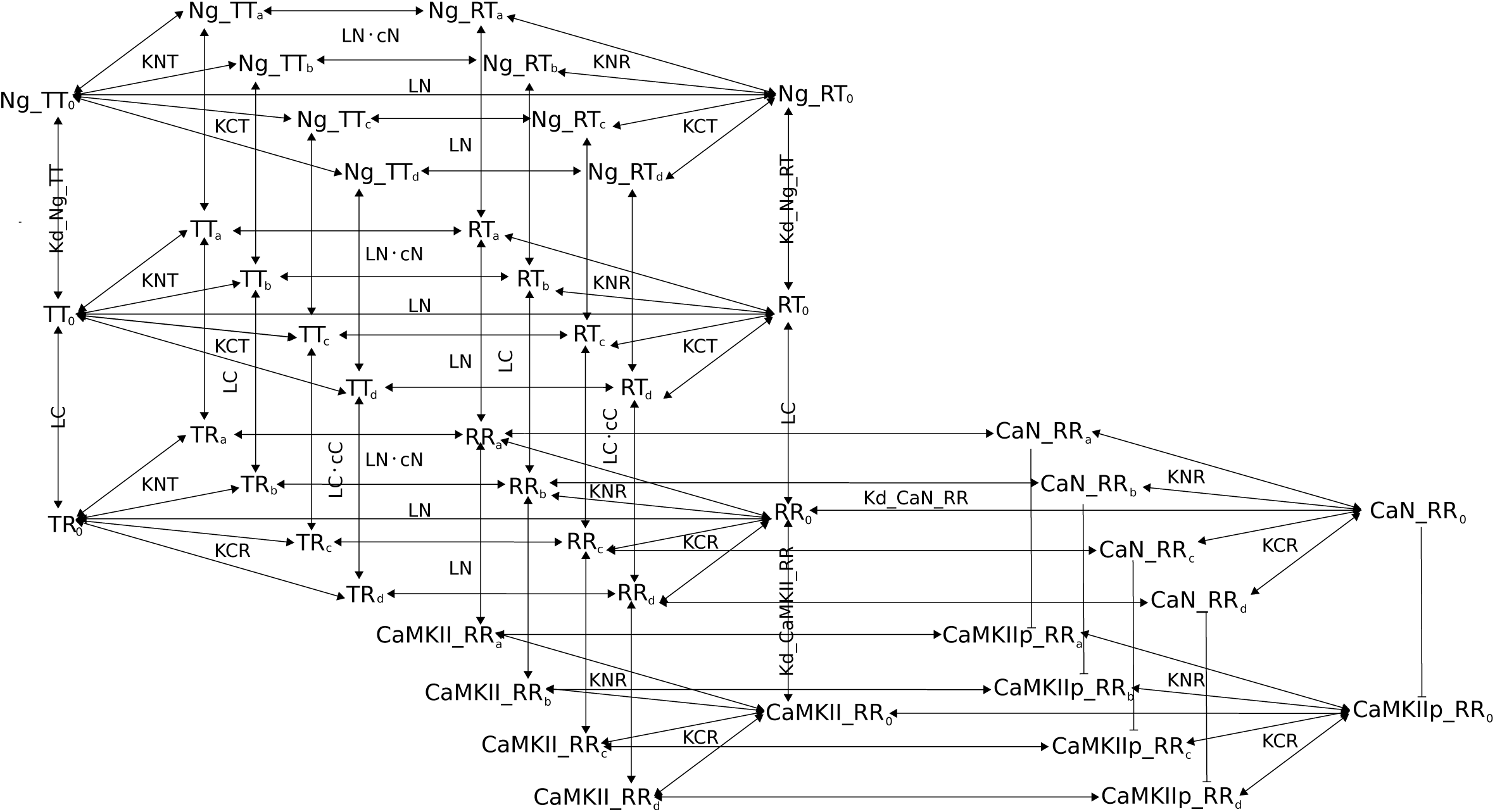
Reaction Diagram involving calmodulin and its binding partners. Binding interactions and state transitions involving calcium, calmodulin and calmodulin-binding proteins. For simplicity, only the binding of the first calcium ion is included. The reactions shown here are applied to all other calcium-bound forms, with any combination of filled calcium-binding sites. a and b represents calcium binding on calmodulin N lobe; c and d represents calcium binding on the C lobe. R indicates open lobes; T closed lobes. Dissociation coefficients for calcium at specific binding sites are specified, as well as state transition constants (ratios between opposite state transition rates). Identical dissociation constants for protein binding calmodulin are only written once. Parameter values are included in Table 1.

First, we modified the hemiconcerted calmodulin model by Lai *et al.* 2015 [25], to allow the binding of calcium ions to affect equally, but in opposite directions, both transitions of calmodulin lobes between open and closed conformations. In other words, calcium binding not only speeds up the T to R state transition, but also slows down the R to T state transition. We also removed the assumption that protein partners can bind to all possible conformations of calmodulin. Rather, we defined these bindings based on the described properties of each specific protein.

**Table 1.** List of parameters used in the model.

| Estimated parameters for the N lobe |  | Estimated parameters for the C lobe |  |
| --- | --- | --- | --- |
| Allosteric parameters: |  |  |  |
| $cN$ | 0.01 | $cC$ | $5 \cdot 10^{-5}$ |
| $LN$ | 299 | $LC$ | 195865 |
| Dissociation constants of calcium from calcium lobes: |  |  |  |
| $KNT$ | $9.38 \cdot 10^{-5} \text{ M}$ | $KCT$ | $9.38 \cdot 10^{-5} \text{ M}$ |
| $KNR$ | $KNT \times cN$ | $KCR$ | $KCT \times cC$ |
| Calcium binding rates: |  |  |  |
| $kon\_N$ | $1.22 \cdot 10^{10} \text{ M}^{-1} \cdot \text{s}^{-1}$ | $kon\_C$ | $1.22 \cdot 10^8 \text{ M}^{-1} \cdot \text{s}^{-1}$ |
| calmodulin state transition rates (0,1,2: number of calcium ions bound to that lobe): |  |  |  |
| $k\_TtoR\_N0$ | $316227 \text{ s}^{-1}$ | $k\_TtoR\_C0$ | $316227 \text{ s}^{-1}$ |
| $k\_TtoR\_N1$ | $k\_TtoR\_N0/\sqrt{cN}$ | $k\_TtoR\_C1$ | $k\_TtoR\_C0/\sqrt{cC}$ |
| $k\_TtoR\_N2$ | $k\_TtoR\_N0/cN$ | $k\_TtoR\_C2$ | $k\_TtoR\_C0/cC$ |
| $k\_RtoT\_N0$ | $k\_TtoR\_N0 \cdot LN$ | $k\_RtoT\_C0$ | $k\_TtoR\_C0 \cdot LC$ |
| $k\_RtoT\_N1$ | $k\_RtoT\_N0 \cdot \sqrt{cN}$ | $k\_RtoT\_C1$ | $k\_RtoT\_C0 \cdot \sqrt{cC}$ |
| $k\_RtoT\_N2$ | $k\_RtoT\_N0 \cdot cN$ | $k\_R2T\_C2$ | $k\_R2T\_C0 \cdot cC$ |
| Parameters relating to calmodulin-binding substrates: |  |  |  |
| $k\_CaMKII\_autop$ | adjusted\_coefficient $\times 6.3 \text{ s}^{-1}$ | This is adjusted at each time step during simulation. Base rate is from [58]. | |
| $koff\_CaMKII\_RR$ | $1.1 \text{ s}^{-1}$ | CaMKII's dissociation rate from active calmodulin [53]. | |
| $kon\_CaMKII\_RR$ | $2.87 \cdot 10^6 \text{ M}^{-1} \cdot \text{s}^{-1}$ | Estimated CaMKII's binding rate to active calmodulin | |
| $koff\_CaMKIIp\_RR$ | $8.7 \cdot 10^{-5} \text{ s}^{-1}$ | Thr286-phospho-CaMKII's dissociation rate from active calmodulin [53] | |
| $kon\_CaMKIIp\_RR$ | $3.85 \cdot 10^7 \text{ M}^{-1} \cdot \text{s}^{-1}$ | Estimated Thr286-phospho-CaMKII's binding rate to active calmodulin | |
| $kon\_CaN\_Ca1, 2, 3, 4$ | $2 \cdot 10^7 \text{ M}^{-1} \cdot \text{s}^{-1}$ | Calcineurin B's binding rate to Ca [10] | |
| $koff\_CaN\_Ca$ | $0.0092 \text{ s}^{-1}$ | [10] | |
| $koff\_CaN\_Ca\_Ca$ | $0.0312 \text{ s}^{-1}$ | [10] | |
| $koff\_CaN\_Ca2\_Ca$ | $0.352 \text{ s}^{-1}$ | [10] | |
| $koff\_CaN\_Ca3\_Ca$ | $0.9 \text{ s}^{-1}$ | [10] | |
| $kon\_CaN\_RR$ | $2.3 \cdot 10^7 \text{ M}^{-1} \cdot \text{s}^{-1}$ | Estimated Calcineurin's binding rate to activated calmodulin | |
| $Kd\_CaN\_RR$ | $3.26 \cdot 10^{-9} \text{ M}$ | Estimated Calcineurin's dissociation constant from activated calmodulin | |
| $kon\_Ng\_RT$<br>$kon\_Ng\_TT$ | $1 \cdot 10^{14} \text{ M}^{-1} \cdot \text{s}^{-1}$ | Estimated neurogranin's binding rate to calmodulin when C lobe is closed | |
| $Kd\_Ng\_RT$<br>$Kd\_Ng\_TT$ | $1.2 \cdot 10^{-6} \text{ M}$ | Estimated neurogranin's dissociation constant from calmodulin, when C lobe is closed | |
| $kcat\_CaN\_CaMKIIp$ | $0.8 \text{ s}^{-1}$ | Calcineurin's catalytic efficiency on dephosphorylating CaMKII<br>Used in this paper | |
| $Km\_CaN\_CaMKIIp$ | $2 \cdot 10^{-5} \text{ M}$ | Calcineurin's Michaelis-Menten constant on dephosphorylating CaMKII<br>Used in this paper | |

For parameter estimations, please see method and supplementary sections. R: open; T: closed. When combined together, the first letter indicates the state of calmodulin's N lobe while the second one indicates the state of the C lobe. Since $kon$ , $koff$ , and $Kd$ are linked ( $Kd = koff/kon$ ), only two of these three are displayed, depending on which ones were actually estimated. For parameters concerning sequential dissociation of $\text{Ca}^{2+}$ ions from CaN, the first "Ca" indicates the number of $\text{Ca}^{2+}$ ions bound to CaN, the second "Ca" shows the one dissociating from CaN.

CaMKII monomer binds to calmodulin when both lobes are in open states (namely the “RR” conformation). This binding exposes the kinase domain of CaMKII monomer, allowing it to phosphorylate and to be phosphorylated by its neighbour monomers within the same hexamer ring, if they are also in active conformation [45]. We adapted and improved the approach used by Li *et al.* 2012 [10] to compute the probability of having an adjacent active monomer, at each time step, based on the proportion of active monomers in the whole system (calmodulin bound and/or phosphorylated). For more details, see the section CaMKII autophosphorylation below. We then used this probability to adjust the global autophosphorylation rate of CaMKII monomers. Once phosphorylated, CaMKII monomers have a higher affinity for calmodulin than their non-phosphorylated counterparts, and remain active even upon calmodulin dissociation [46]. CaMKII monomers are dephosphorylated by Protein Phosphatase 1 (PP1). As CaN activates PP1 “linearly” through dephosphorylation of Thr34-phospho-DARPP-32, which then releases PP1 inhibition, we simply modelled a direct dephosphorylation of CaMKII monomers by CaN using Henry-Michaelis-Menten kinetics with total active CaN as the enzyme concentration.

CaN is a heterodimer containing a regulatory subunit (CaNB), and a catalytic subunit (CaNA) [47]. In our model, the sequential binding of four calcium ions to CaNB is required before CaNA can bind calmodulin, thereafter becoming active [48, 49]. CaNA binds to calmodulin when both calmodulin lobes are in the open state (“RR”).

As Ng interacts mostly with the C lobe, in this model we considered that it bound to calmodulin only when the C lobe was in the closed state, regardless of the N lobe conformation (“RT” and “TT”). As a consequence, the association of calmodulin with Ng does not prevent the transitions of the N lobe between T and R states. Moreover, due to the lack of relevant experimental data, we assumed that the binding of Ng on the C lobe did not exert any allosteric effect on those transitions.

To create a basal level of calcium (0.08 · 10^−6^ M), we implemented reactions to create constant influx and removal of ions, by mimicking a passive calcium input channel and a concentration-dependent calcium pump. Calcium input was encoded as a train of calcium spikes at varied frequencies. Each calcium spike was generated by adding a zero-order calcium creation during 8-millisecond (mimicking the opening of calcium channels), that elevated the free calcium concentration transiently to 0.08 · 10^−6^ M. The spike’s half-life time was about 30 ms, corresponding to the experimental observation [50]. For each free calcium spike, a total of 1926 calcium ions were injected in the model over a 8 ms period.

### Estimation of equilibrium constants for calmodulin and calcium

The function of allosteric proteins is characterised by two types of parameters. The thermal equilibrium is quantified by *L*, the concentration ratio between the different states in the absence of any allosteric effector. In our case, calmodulin lobes exist under two states, closed (T), and open (R) and for each lobe, we have an allosteric constant *L* = [*T*]*/*[*R*]. Extremely large or extremely small *L* indicates a system for which the equilibrium is strongly biased towards a given conformation. In addition, for each allosteric effector, the ratio of affinities for the two conformations, dubbed *c* for calcium or *e* for neurogranin, indicates if a ligand favours one conformation or the other.

We estimated the allosteric parameters for both N and C lobes by using experimental data obtained predominantly from full-length CaM, where intrinsic phenylalanine and tyrosine fluorescence were used for monitoring calcium saturation. We used three sets of steady-state experimental data: 1) calmodulin titrating CaMKII peptide with a saturating amount of calcium ions [6], 2) calcium saturation curves of the C lobe in the full length calmodulin either without targets or in the presence of CaMKII peptide or full-length Ng [6, 26], 3) calcium saturation curve of the N lobe, in full-length calmodulin, without targets [6]. As the experimental data concerning N lobe calmodulin is relatively scarce, we also used a calcium titration curve of truncated N lobe of calmodulin, that was locked in closed conformation [51].

To fully characterise the interactions between calmodulin and calcium, we therefore have to estimate: 1) the binding affinities of calcium to the T state of calmodulin for sites on both lobes. Following Lai *et al.* [25], we further hypothesized that within each calmodulin lobe, the affinities of the two calcium binding sites were the same and we had only two affinities to estimate (*KNT* and *KCT*), 2) the ratio between calcium affinity for the R and T states for both lobes (*cN* and *cC*) which we assumed to be equal for the two binding sites within each lobe as in Lai *et al.* 2015 [25], 3) the ratio of T state calmodulin to R state in the absence of calcium for both lobes (*LN* and *LC*), and 4) the affinity of calmodulin to the CaMKII peptide (*Kd*_CaMKIIpep_*RR*) and to Ng (*Kd*_Ng_*TT* and *Kd*_Ng_*RT*).

As for the full-length CaMKII, the CaMKII peptide used in the published experimental datasets binds to calmodulin when both lobes are in R state. Ng only interacts with calmodulin’s C lobe and, as explained in the model structure section, *Kd*_Ng_*RT* = *Kd*_Ng_*TT*, i.e. Ng binding does not affect the state transitions of the N lobe and *vice versa*. The parameters remaining to be estimated are therefore reduced to: *KNT*, *KCT*, *cN*, *cC, LN*, *LC, Kd* CaMKIIpep *RR*, and *Kd*_Ng_*TT*.

Even with the assumptions described above, the amount of parameters to estimate is large and correlations may arise between them. Moreover the experimental conditions used in these estimation procedures are highly diverse. Thus, we proceeded in several stages.

Firstly, using calcium titration experiments of truncated N lobe locked in the closed conformation by disulfide crosslinks [51], we estimated the affinity between calcium and the N lobe in the T state to be *KNT* = 9.38 · 10^−5^ M (S1 Fig.a and b).

Using experimental measurements where calmodulin titrates CaMKII peptide in the presence of saturating amount of calcium ions [6], we then estimated the affinity between calmodulin and CaMKII peptide (*Kd*_CaMKIIpep_*RR*). Because of the high calcium concentration we assumed that almost all calmodulin molecules were in the R state. This gave a value of *Kd*_CaMKIIpep_*RR* = 5.6 · 10^−9^ M (S1 Fig.c and d). This only provides the upper bound for the dissociation constant between CaMKII peptide and RR calmodulin, as not all calmodulin molecules are in the RR state, even with the highest calcium concentration. In fact, this value was reduced to 3.2 · 10^−10^ M during subsequent stages of parameter estimations.

Estimating parameter values requires sampling values within given ranges. We calculated the boundary values of KCT and cC by assuming that the observed calcium saturation levels of the C lobe, in the presence either of CaMKII peptide or of Ng [6, 26], reflected the calcium binding affinities when all C lobes were locked in the R or T state respectively, resulting in *cC*^*I*^ = 0.0011 (*cC* = *KCR/KCT*) and *KCT* ^*I*^ = 1.3 *·* 10^−5^ M (S1 Fig.e-g). This value of *cC*^*I*^ was used as the upper bound for estimating the real *cC*, as in reality the CaMKII peptide is not capable of locking all C lobes in the R state. Similarly, *KCT* ^*I*^ was used as the lower bound for *KCT*, since not all C lobes are locked in the T state by Ng, i.e. the actual affinity of the T state for calcium is lower.

Since *KCT* had the same order of magnitude as our estimated *KNT*, and calcium affinity for the C lobe should be higher than for N lobe, the range for the real *KCT* value was narrow. Thus, to further reduce the amount of parameters to be estimated, we simply assumed that, in the closed state, both C and N lobe shared the same calcium binding affinities, that is *KCT* = *KNT* = 9.38 · 10^−5^ M.

Finally, we estimated the remaining parameters: *cC, cN*, *LC, LN* and *Kd Ng TT* together using the full calmodulin model with steady state calcium titration curves [6, 26] and the boundary values mentioned above. As shown in S2 Fig., all parameters except *cC* and *LC* are identifiable. Fitting results are illustrated in S3 Fig.. As Lai *et al.* [25], we found *cC* and *LC* highly correlated. Fitted their logarithms with a linear regression model provided a slope of -1.9 (S4 Fig.). We therefore further refined these two parameters while estimating the rate constants.

### Estimations of kinetic rate constants

We estimated the affinity between open calmodulin (conformation “RR”) and calcineurin by using the CaN dose-response to calmodulin, in the presence of saturating calcium concentration [52]. We also estimated the association constant between the two proteins using stopped-flow kinetic data, where mutant calmodulin was labelled by Acrylodan (*CaM* (*C*75)_*ACR*_) [52]. The estimated affinity was *Kd*_CaN_*RR* = 3.2 · 10^−9^ M, with an association constant *kon*_CaN = 2.3 10^7^ M^-1^·s^-1^ (S5 Fig.).

We directly used the dissociation constants of CaMKII and phospho-CaMKII from calmodulin taken from the literature (*koff*_CaMKII_*RR* = 1.1 s^-1^; *koff*_CaMKIIp_*RR* = 8.7 · 10^−5^ s^-1^) [53], and therefore only had to estimate their association rate constants. Based on previous published works [23, 26], we hypothesized that calcium binding to the N lobe was 100 times faster than to the C lobe, (*kon_N* = 100 *× kon_C*), and these did not depend on the conformation of the lobes, therefore reducing the estimation of 8 unknown association rates to only 1, *kon CT*. Finally, we assumed that the base state-transition rates were the same for the two lobes, *k_TtoRC*0 = *k_TtoRN* 0. The specific state transition rates, for all liganded populations, depend on the number of calcium bound to the calmodulin lobe, the protein partner bound, and the allosteric parameters estimated for this lobe. We also made use of the relationship between *cC* and *LC* described above, and only had to estimate *cC*.

We used stopped-flow fluorometry measurements of quin-2 fluorescence increase, after addition of calcium-saturated calmodulin mix, in the presence of either CaMKII or phospho-CaMKII [31], and measurements of native tyrosine fluorescence decrease in the calmodulin C-terminal in the presence of EGTA, with or without neurogranin [26].

Most of the parameters estimated at this stage were identifiable, except the association constant between Ng and calmodulin *kon_Ng*, as well as the base state-transition rate *k_TtoRC*0 (S6-7 Figs.). Thus, we chose the highest reasonable association constant for Ng and calmodulin (*kon*_Ng = 10^14^ M^-1^·s^-1^) and an average value for the transition rate among the range of values that all reached best fit (*k_T* 2*RC*0 = 316227). The fitted results are illustrated in S8 Fig..

### CaMKII autophosphorylation

During simulations, we actively adjusted the global autophosphorylation rate of CaMKII monomers based on the total amount of active monomers, that are the phosphorylated and/or calmodulin bound monomers, and the likelihood of them having adjacent active monomers in pseudo hexamer rings, an updated approach from our previous work [10].

Briefly speaking, there are 7 types of hexamers, containing 0 to 6 active monomers. For each type, the possibilities of location for the active monomers are limited and we can easily compute the probability for an active monomer to have a neighbour also active. The main aspect of the procedure is to approximate the fraction of the different types of hexamers when the total number of active monomers is updated from the simulation. We achieved this by running, for every 1 percent increase of active monomers, 1000 independent random distributions of the active monomers to hexamers. For every increase of active monomers, we then computed the average occurrences of each type of hexamers, and transformed them into the probability for a monomer to be distributed in a specific type of hexamer. We then multiplied the distributions of monomers with their corresponding probability of adjacent active neighbours. Finally, the summation of the above were used as the overall probability and the chance of having an active neighbour monomer for each percent increase in total active monomers. We fitted this data with a 5^th^ order polynomial function and embedded it into the model to adjust the autophosphorylation rate of CaMKII monomer (S9 Fig.).

### Numerical simulations and analysis of results

All simulations, including timecourses with calcium spikes were performed with COPASI [54], using the LSODA solver [55]. Parameter estimations were conducted using the SBPIPE package [56], combined with the “Particle swarm” optimization method (2000 iterations with a swam size of 50). Parameters used for simulation of the model are listed in Table 1.

The frequency of calcium spikes was directly encoded in the model together with the number of spikes, in order to control the total duration of calcium inputs. We ran the simulations on a computing cluster, using Python’s *ElementTree* package to automatically modify the frequency parameter in CopasiML files.

Each simulation started with a 5000 s equilibrium - with output interval size set at 1 s, followed by the simulation of 300 calcium spikes at frequencies ranging from 0.1 Hz to 100 Hz, with output interval size set at 0.0001 s. We recorded the evolution of all protein active forms during 3000 s.

In order to evaluate CaMKII and CaN’s effects on synaptic plasticity, we measured the “activated area” [10], which was obtained by integrating, over the 3000 s, the product of their relative activations (concentration of active proteins over total concentration) above basal activities (substracting basal levels), by their catalytic constants on AMPA receptor GluR1 subunit. The BCM-like curve [57], useful to characterise bidirectional plasticity, was obtained by subtracting the activated area of CaN from that of CaMKII for each calcium spike frequencies.

## Results

### Neurogranin sequesters closed-state calmodulin

We firstly studied how calmodulin (CaM) binding proteins can affect calmodulin’s apparent affinity for calcium at steady states. As shown in Fig. 2, and in accord with previous experiments [6, 26] and modelling [25], the presence of Ng shifts the calcium-saturation curve of calmodulin’s C lobe towards higher calcium concentration range, reducing the apparent affinity for calcium. This indicates that Ng traps the C lobe in the closed, low-affinity, conformational state, and therefore hinders calcium binding, in particular at low calcium concentrations. Conversely, CaMKII and CaN promote calcium binding to the C lobe, increasing the apparent affinity, through their preferential binding to the open, high-affinity conformation (of both lobes).

**Fig 2.**
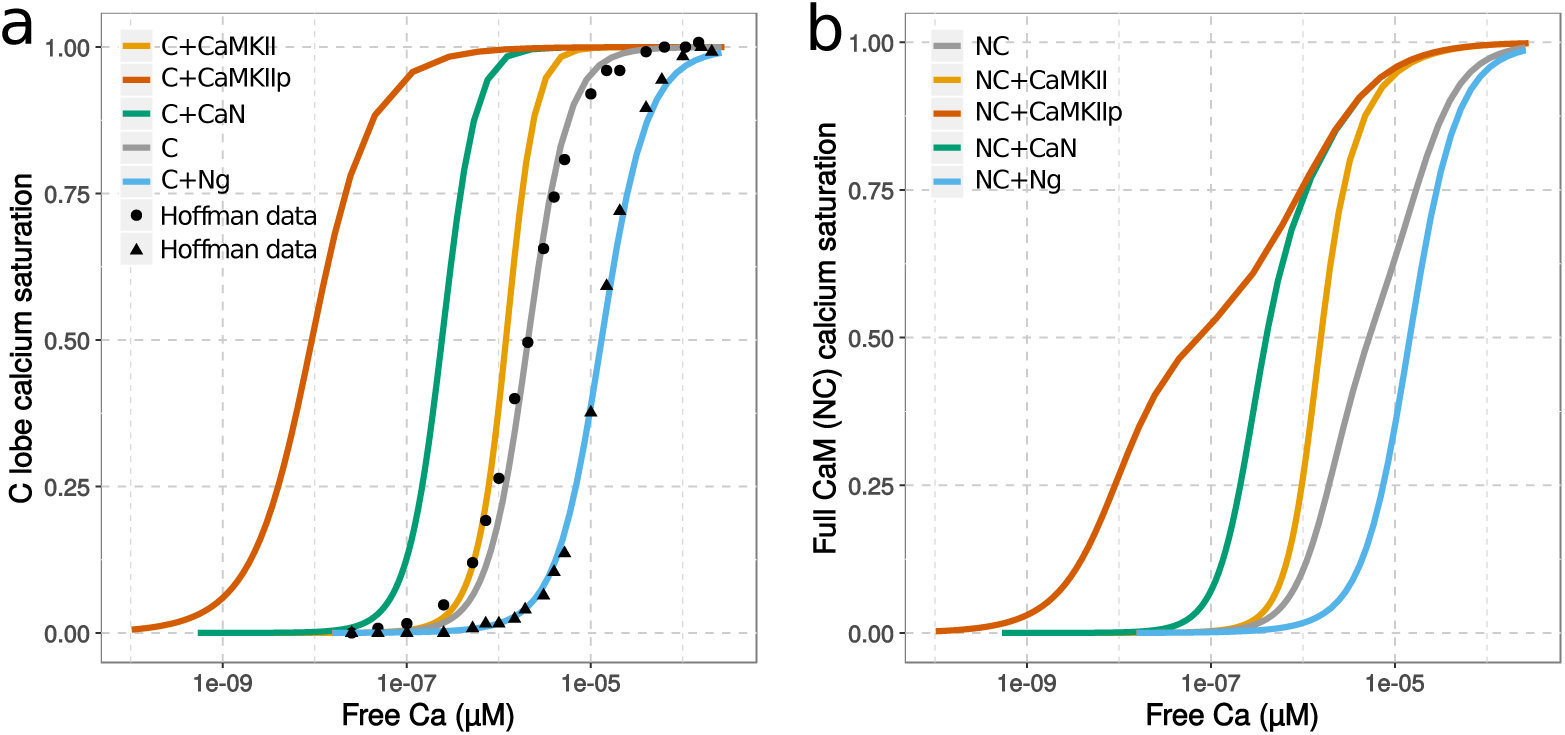
Saturation curves of calmodulin by calcium. Values were obtained by scanning a range of initial calcium concentrations and running simulations until steady-state is reached, using initial conditions described in Hoffman et al. 2014 [26]. a) shows only the saturation of C lobe (within the context of the entire calmodulin) while shows the whole calmodulin, without any substrates (grey), with neurogranin (blue), non-phosphorylated CaMKII (orange), Calcineurin (green) or Thr286-phospho-CaMKII (red). The x-axis shows the free concentration of calcium, and not the initial concentration used for the scan. [CaM] = 5 µM; [Ng/CaMKII/CaMKIIp/CaN] = 50 µM. Solid line: simulation results; dots: experimental observations.

Calcium saturation curves for full calmodulin show similar trends as for the C lobe (Fig. 2), but they also indicate calmodulin’s lobe differences in terms of calcium binding. Based on the allosteric parameters we estimated, the N lobe is more flexible and has greater potential to undergo spontaneous conformational transitions than the C lobe (*LN* closer to 1 than *LC*). However, the binding of calcium has less influence on the conformation of the N lobe, because of the the smaller difference between the affinities for calcium towards the two conformations (*cN* closer to 1 than *cC*) in comparison with C lobe. The saturation curve of full calmodulin in the absence of protein binding partners is slightly stretched, showing a larger contribution of the C lobe at lower calcium concentrations. As in our model Ng only interacts with C lobe, it bears little effect on N lobe’s calcium binding, and the shift is mostly seen at lower calcium concentrations. Whereas CaMKII and CaN are wrapped around by both N and C lobes, and their presences greatly enhance, not only C lobe but also N lobe’s calcium binding. However, N lobe’s affinity to calcium is less increased by the conformational change to open state. Although having the highest affinity towards calmodulin, phospho-CaMKII cannot shift N lobe’s affinity much compared to non-phospho CaMKII.

In line with the findings above, timecourse simulations show that Ng speeds up calcium dissociation from calmodulin’s C lobe (Fig. 3), in agreement with stopped-flow fluorometry experimental observations [26]. Taking steady-state and kinetic results together, these indicate that Ng not only decreases calcium binding to calmodulin at steady states, but also dynamically increases calcium dissociation from calmodulin when free calcium concentration is decreased. Both observations arise from the preferential binding of neurogranin to a conformation of calmodulin’s C lobe with low affinity for calcium.

**Fig 3.**
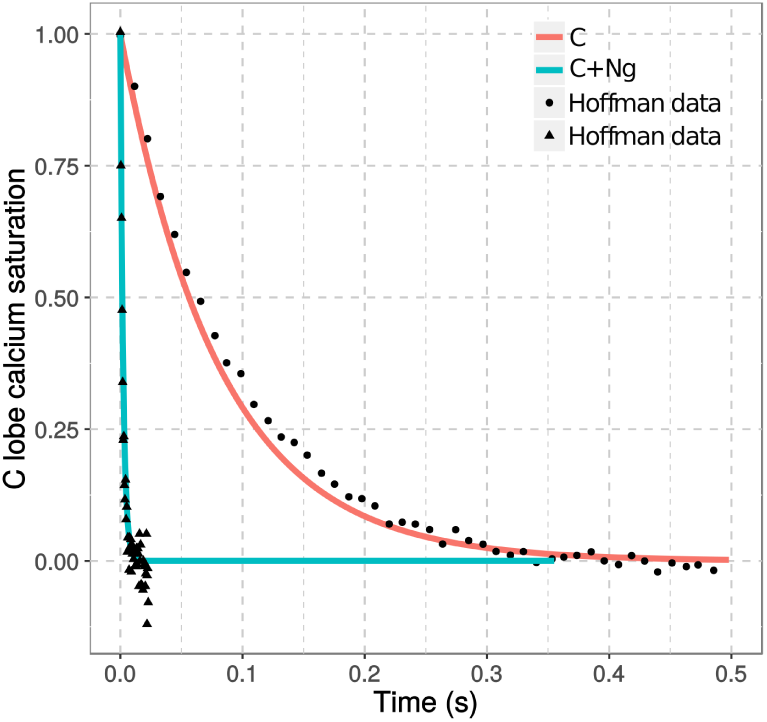
Kinetics of calcium dissociation from calmodulin. Calcium dissociation from the C lobe of calmodulin was measured upon mixing calcium chelator EGTA (1 · 10^−2^ M) after model simulation reaching steady states with pre-mixed calmodulin (10 µM) and calcium (100 µM) with (cyan) or without (salmon) neurogranin (50 µM). Solid line: simulation results; dots: experimental observations [26].

### Neurogranin shifts calmodulin activation towards high-frequency calcium spikes

Post-synaptic events trigger calcium spikes that can increase free calcium concentration transiently to sub-micromolar levels in a few milliseconds, subsequently declining in about 200 milliseconds, due to calcium pumps and calcium binding proteins [50]. Therefore, rather than looking at the responses of calmodulin to calcium steady-states, we must look at its responses to calcium spike frequencies (Fig. 4).

**Fig 4.**
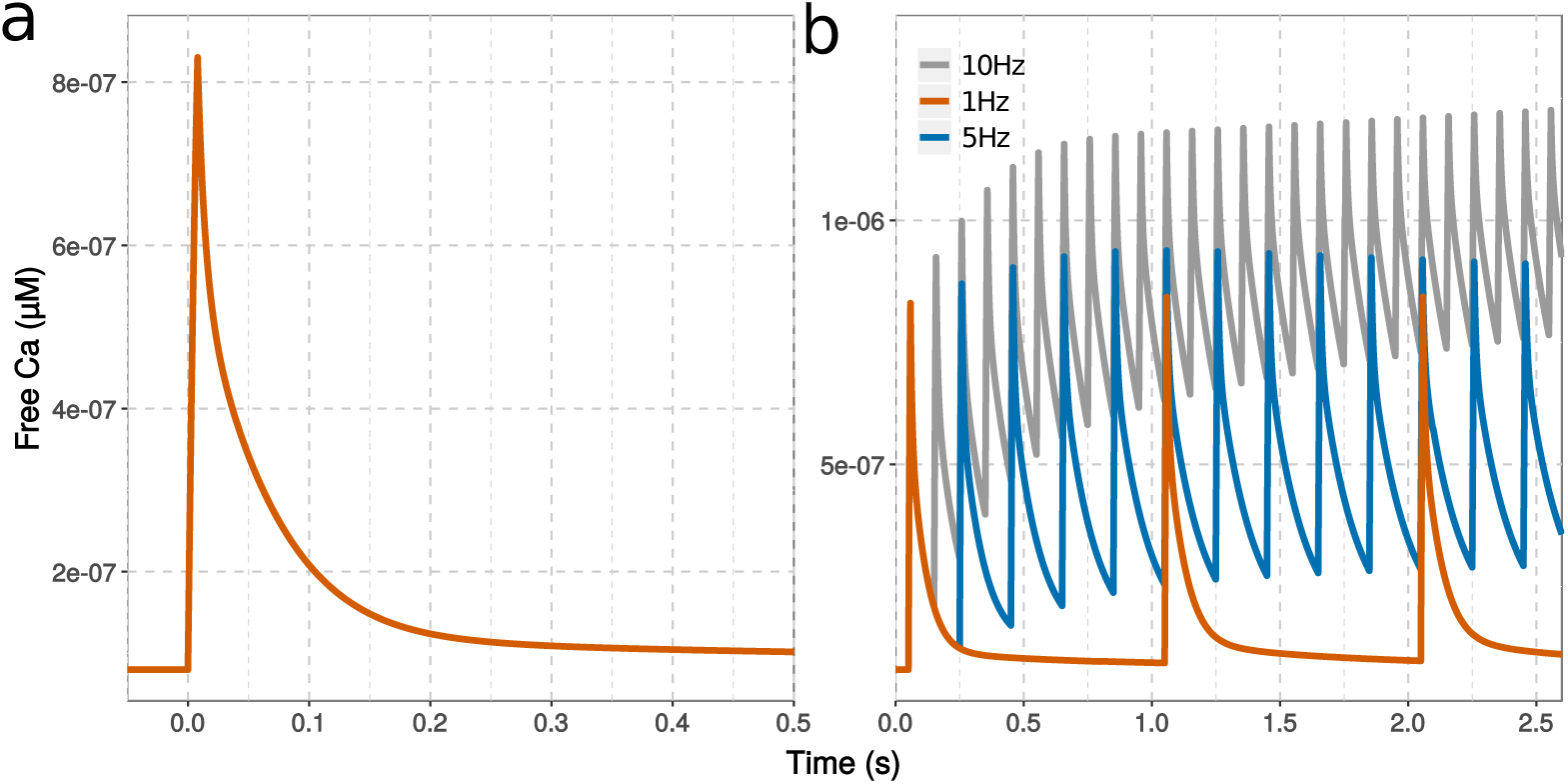
Free calcium concentration at various input frequencies. Intracellular free calcium elevation simulated with a single calcium input (a) or a train of inputs (b) at 1 Hz (red), 5 Hz (blue) and 10 Hz (grey). Each calcium spike represent the addition of 1926 molecules over 8 ms

We first simulated the calmodulin model, without protein partners, using a fixed total calcium input applied at varied spike frequencies. The results are in line with the behaviour of our previous fully concerted model [10], and show that both calmodulin lobes respond differently to different calcium spike frequencies. They stay in the open state for about 10 s at high frequencies, even when the elevation of calcium concentration is much shorter, which is not shown in the steady state analysis (Fig. 5). However, the N and C lobes display different sensitivities towards calcium spike frequencies. The C lobe opens at lower frequencies, and for longer durations, whereas the N lobe requires much higher spike frequencies to switch to the open conformation. The openings of both lobes follows calcium binding tightly.

**Fig 5.**
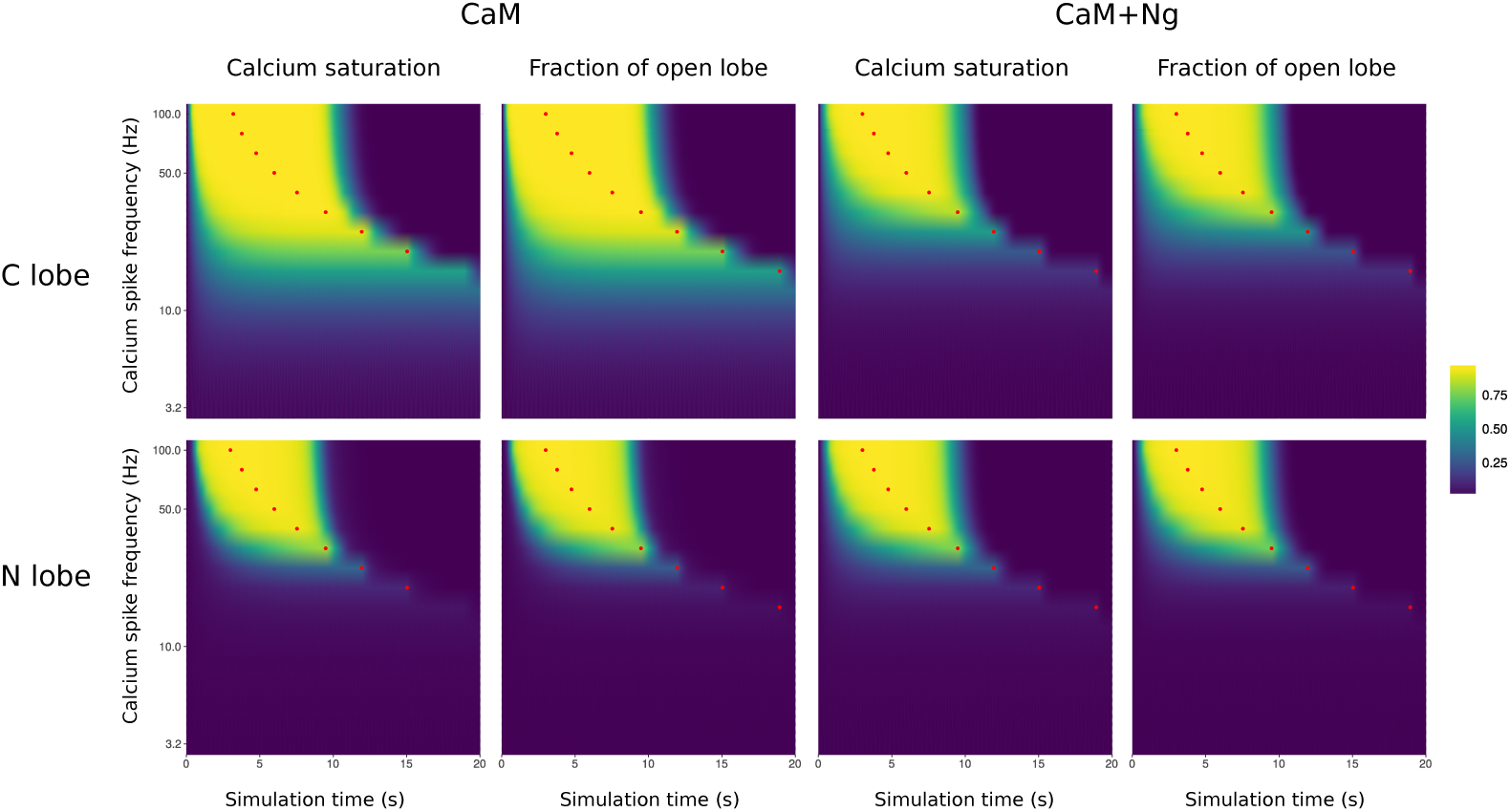
Calcium binding and calmodulin conformational changes in response to calcium input frequencies. Calcium binding calmodulin was followed during simulations using trains of 300 spikes at different frequencies. Plots show calcium saturation levels at the C and N lobe and the corresponding calmodulin conformational changes, with calmodulin on its own, or in the presence of neurogranin. Red dots represent the end of calcium stimulations. [CaM]_tot_ = 40 µM, [Ng]_tot_ = 40 µM, basal [Ca] = 0.08 µM.

When Ng was added to the model, the range of frequencies able to open most C lobes was narrowed towards high values (Fig. 5). The duration of the activation was also shortened. Unsurprisingly, no changes were observed for the N lobe (Fig. 5). Overall, the opening patterns of the C lobe in presence of Ng resembles those of the N lobe, restricting the opening to high calcium input frequencies.

To better quantify calmodulin’s responses to the frequencies of calcium input, we calculated the Area Under the Curve (AUC), that is the integral of the increased/decreased fraction of calmodulin in the open/closed state (above/below basal level), over the entire simulation (see methods). We calculated these AUCs for all the possible conformations of calmodulin lobes: RR,RT,TR and TT (where the first letter represents the N lobe and the second represents the C lobe. We then plotted them against calcium spike frequencies (Fig. 6).

**Fig 6.**
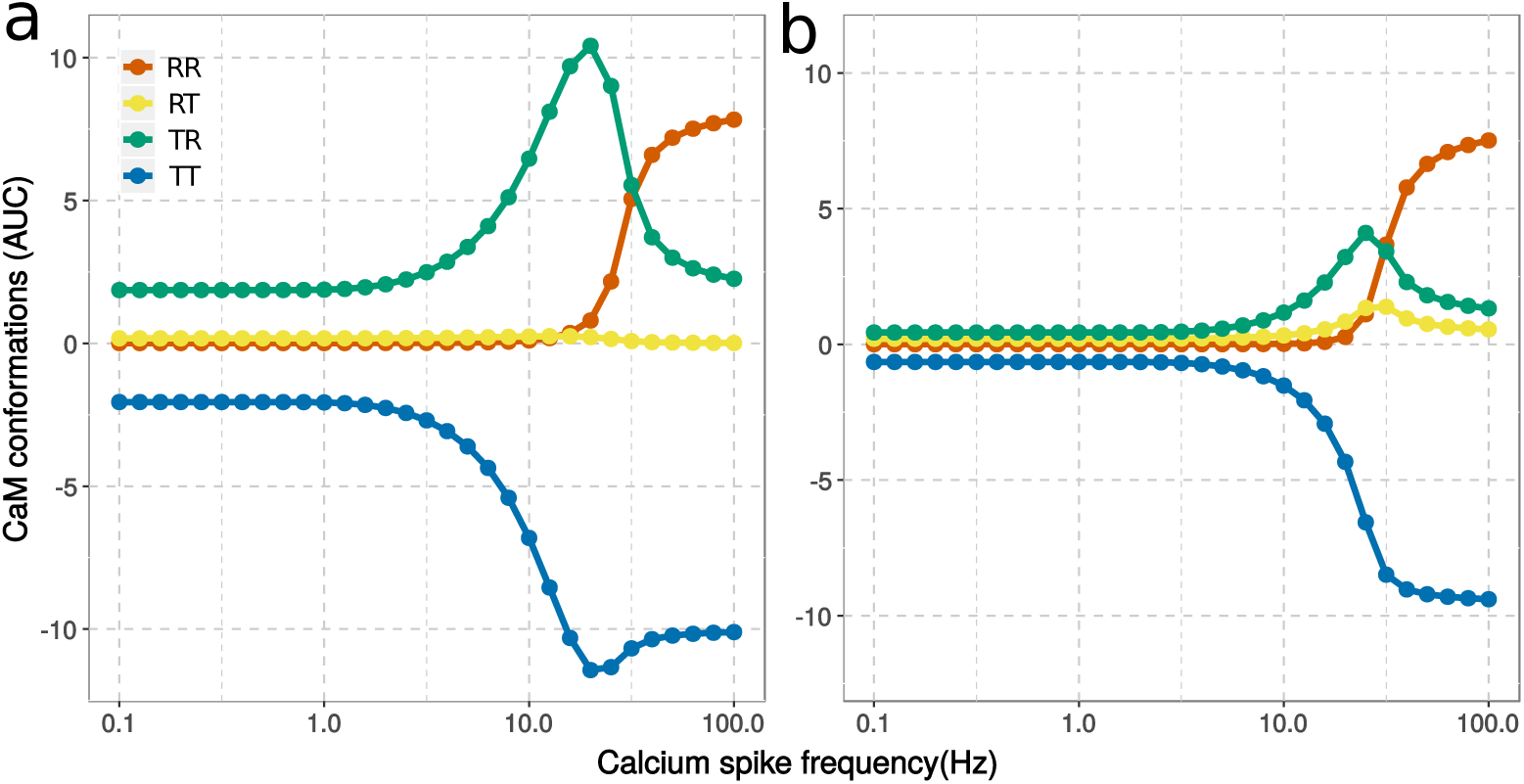
AUCs of calmodulin conformations in response to calcium input frequencies. Conformational changes of calmodulin (40 µM) without neurogranin(a) or with neurogranin ([Ng]_tot_ = 40 µM) (b). Basal activities were subtracted before AUCs were calculated.

Without Ng, Calmodulin’s C lobe is very sensitive to calcium spikes at low frequencies. The conformation TR increases above its basal level (AUC*>*0, as basal activity has been subtracted.) by calcium stimulation as low as 0.1 Hz (Fig. 6a). The conformation TR increases steeply around 3 Hz, peaks at 20 Hz, then declines while N lobe opens (the increase of RR conformation). There is almost one order of magnitude between the frequencies at which the C and N lobe opens prominently (Fig. 6a).

Neurogranin not only suppresses the C lobe opening in response to low frequencies, but also shifts it towards higher frequencies. At about 10 Hz, both TR and RT state start to increase and peak around 25 Hz, indicating the opening of both C and N lobes within different populations of calmodulin molecules (Fig. 6b). In fact, without Ng, the N lobe does not open on its own (Fig. 6a). At these intermediate frequencies, Ng facilitates the opening of the N lobe at the cost of reduced open C lobe. At high spike frequencies, the opening of both lobes (RR state) follows almost the same profile as when Ng is absent, and reaches the same activity level at 100 Hz regardless of Ng’s presence (Fig. 6b).

Therefore, neurogranin does not prevent calmodulin activation by calcium spikes. Instead, neurogranin synchronizes both lobes to respond towards higher calcium spike frequencies, hence narrowing the frequency range at which calmodulin responses.

### Neurogranin facilitates CaMKII activation at high calcium spike frequencies

We then investigated neurogranin’s role on regulating the activations of CaMKII and CaN by open calmodulin. We firstly added only CaMKII and CaN in the model (without Ng), as well as inhibition of CaMKII autophosphorylation by CaN (See Materials and Methods). As above, we simulated trains of calcium spikes at various frequencies but resulting in a fixed additional amount of calcium ions. In order to compute their activity on phosphorylation of AMPA receptor subunit GluR1, we multiplied the fraction of active CaMKII and CaN by their respective catalytic constants (kcat).

Fig. 7 shows example timecourses of protein activity changes following 300 calcium-spike stimulations at 10 Hz or 30 Hz. In the absence of Ng, basal calcium level opens a small proportion of calmodulin, resulting in the basal CaN activity that is higher than CaMKII. At 10 Hz calcium stimulation, the availability of more open calmodulin allows a fast and strong elevation of CaN, companied by a gentler raise of CaMKII activity, reaching to the same levels (Fig. 7a). At 30 Hz, the acute elevation of calcium gives rises to sharp and prominent increases of both CaMKII and CaN, and CaMKII overcomes CaN. However, the activation of CaMKII decays very fast to its resting level, potentially due to the slower deactivation and high basal activity of CaN, which prevents the onset of CaMKII autophosphorylation (Fig. 7b).

**Fig 7.**
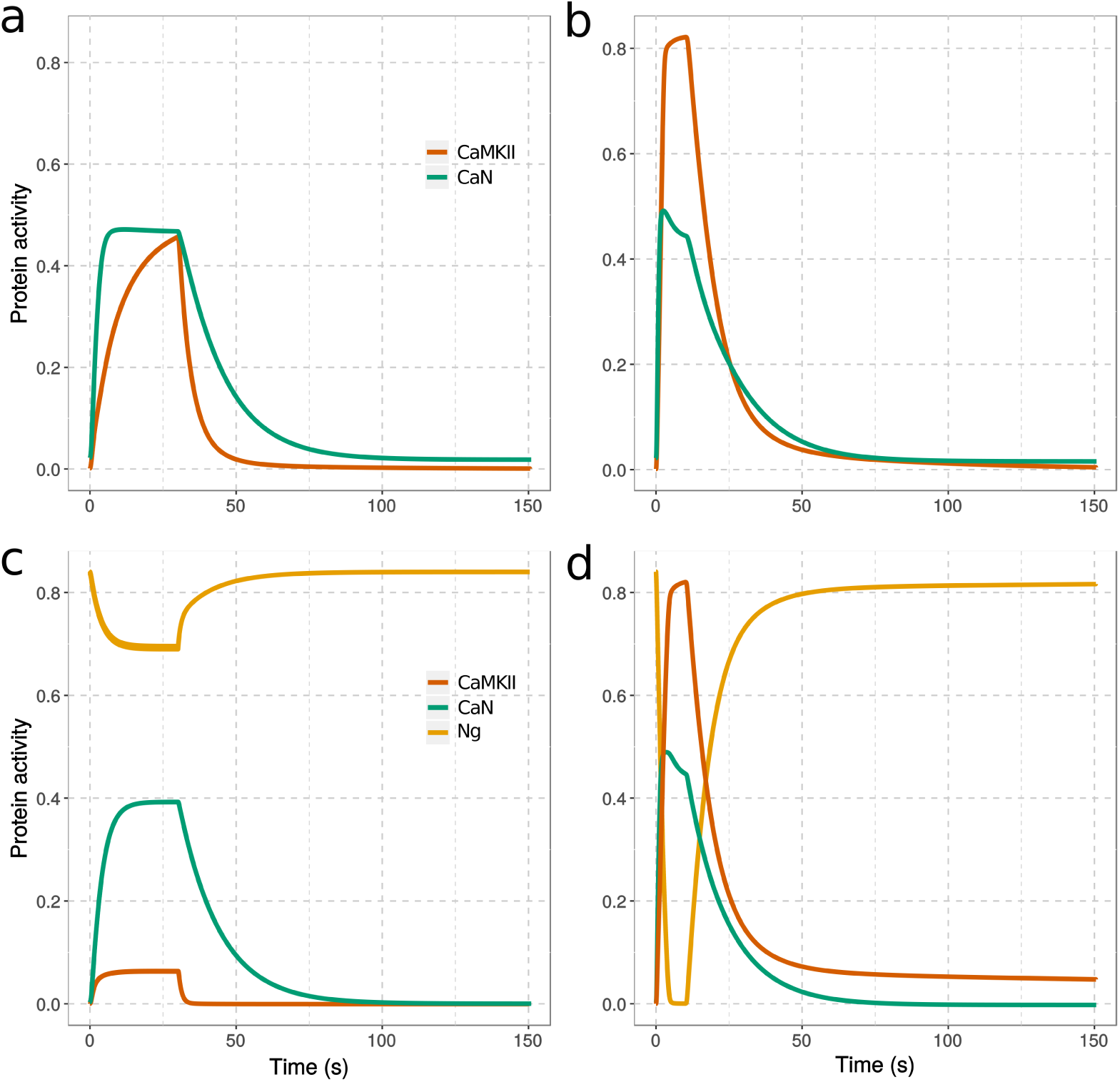
Change of protein activity in response to calcium spikes. Simulations in absence (a,b) or presence (c,d) of neurogranin, from equilibrium and during stimulations by 300 calcium spikes at 10 Hz (a,c) and 30 Hz (b,d). The protein activities were defined as follow: fraction of CaMKII monomers bound to CaM and/or phosphorylated multiplied by CaMKII kcat for GluR1, fraction of CaN bound to CaM multiplied by CaNA kcat for GluR1, fraction of Ng bound to CaM. [CaM]_tot_ = 40 µM, [CaMKII]_tot_ = 80 µM, [CaN]_tot_ = 8 µM, and [Ng]_tot_ = 40 µM; kcat_CaMKII_ = 2 s^-1^, kcat_CaN_ = 0.5 s^-1^. All the concentrations of perspective calmodulin binding proteins are assigned according to the protiomic study in the hippocampus CA1 region [59].

When Ng is present together with CaMKII and CaN in the model, there is less basal CaN activity indicating decreased open calmodulin by Ng binding. Calcium inputs at both frequencies force neurogranin to release open calmodulin, which is then able to stimulate the activity of CaMKII and CaN. At 10 Hz, a small proportion of calmodulin is released from neurogranin, inducing a transient but strong activation of CaN, almost resembles the situation when Ng is absent. The noticeable difference lies on the response from CaMKII, which is much weaker and reaches only a fraction of CaN’s one (Fig. 7c). At 30 Hz, neurogranin releases almost all calmodulin it previously bound. This transient but strong calmodulin release robustly increases CaMKII and CaN activity, which are not so different from without Ng. However, the most striking difference is in the way how CaMKII activity decays. A significant proportion of CaMKII monomers remain active for more than 10 minutes, that is far beyond the total duration of calcium stimulation (Fig. 7d and S10 Fig.). Thus, at high spike frequencies, the presence of Ng promotes the autophosphorylation of CaMKII, which activity lasts beyond calcium spikes and calmodulin association.

In most LTP induction protocols, high frequency stimulation is not applied continuously but grouped into discrete bursts. We verified the finding presented above by splitting the 300 calcium spikes at 100 Hz into three 100 Hz, 100-spike bursts, separated by 10-minute intervals [15]. As shown in Fig. 8, we obtained results similar to a setup featuring a continuous 300-spike stimulation (S11 Fig.). When Ng is present (wt) a small proportion of CaMKII monomers can retain their activity for about 10 minutes after the end of each burst, whereas in the absence of Ng, the elevated basal CaN prevents such a sustained CaMKII activation, resulting in more transient CaMKII activity. Interestingly, with a burst stimulation the overall CaMKII activity (integrated over time) is increased by nearly 20% compared with a continuous 300-spike stimulation.

**Fig 8.**
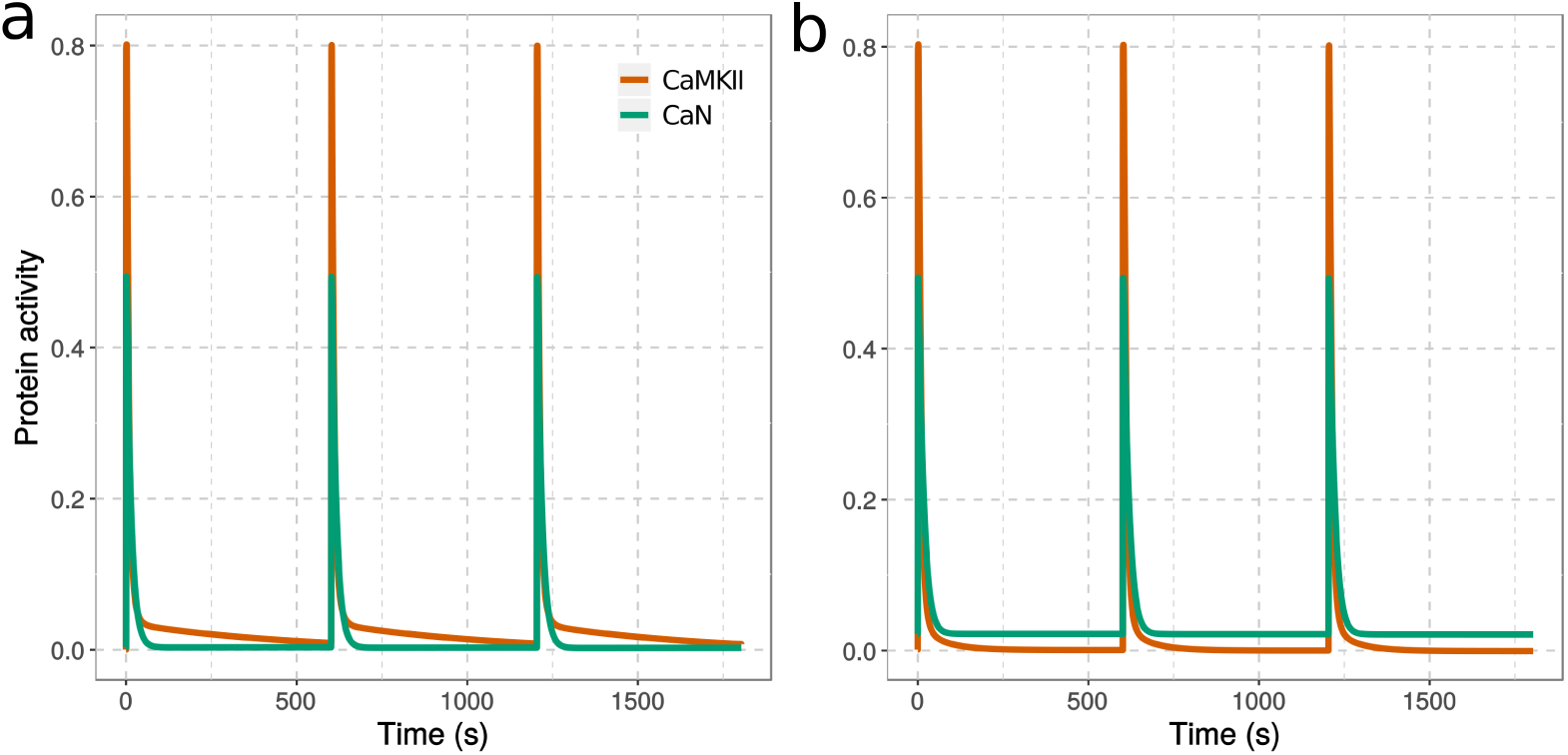
CaMKII and CaN activation in response to tetanic stimulation. Computational models were simulated with (a) or without (b) neurogranin, with 300 calcium spikes organized into three 100-spike discrete bursts at 100 Hz each, and at 10 min interval. Each calcium input was as described in Fig.3. Both protein activities were normalized to their total concentration, then multiplied by their catalytic constant for GluR1. [CaM] = 40 µM, [Ng] = 40 µM, [CaN] = 8 µM, [CaMKII] = 80 µM; *kcat*_CaMKII_ = 2 s^-1^; *kcat*_CaN_ = 0.5^-1^

To represent the combined effect of CaMKII and CaN on GluR1, and thus their impact on synaptic plasticity, we subtracted their activity at each frequency, following a continuous 300-spike stimulation. Negative values indicate that CaN activation overcomes CaMKII, thus facilitating LTD, whereas positive values show that activation of CaMKII overcomes CaN, thereafter facilitating LTP. Fig. 9a shows that in presence of Ng (grey curve), the response to frequencies shows a triphasic curve. At low spike frequencies, CaN is favoured, and this trend increases until around 10 Hz. Above 10 Hz, CaMKII quickly overcomes CaN, reproducing the classical BCM curve. This increase plateaus around 30 Hz, a frequency of stimulation often used to trigger robust LTP. The absence of Ng causes phosphatase activity to be much stronger at low frequencies and less sensitive to frequency increases. The shift towards kinase activity is less sharp, and occurs at lower frequencies, the final plateau representing a much lower kinase activity (Fig. 9a black curve).

**Fig 9.**
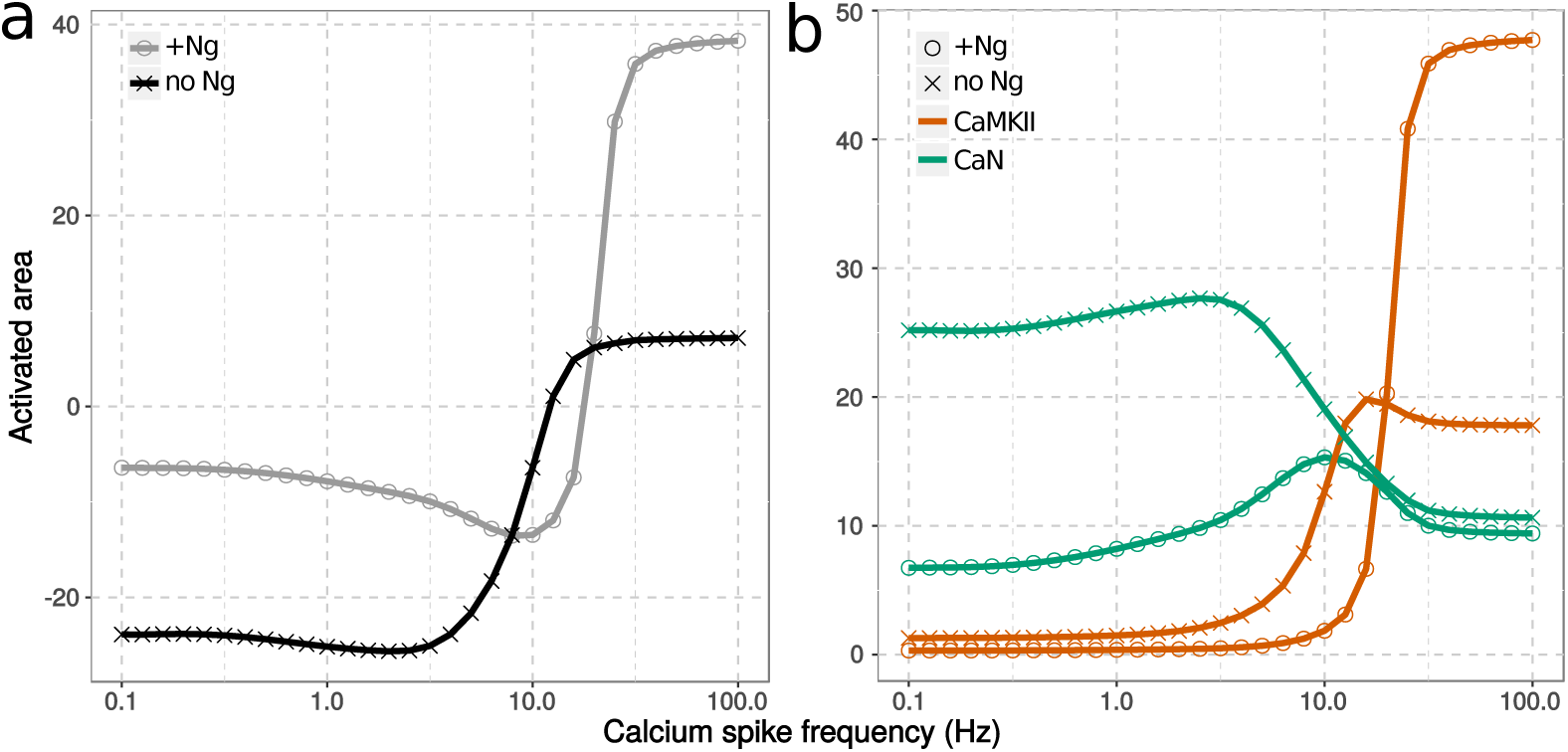
Activity of CaMKII and CaN in response to calcium spike frequencies. Protein activities, as described in Fig 7, were integrated over time. The effect on synaptic plasticity were calculated by subtracting CaN’s activated area from CaMKII’s one (a). Each individual protein’s activated area was also plotted as a function of calcium input frequency (b). Model were stimulated either with (grey circle) or without neurogranin (black cross). All protein concentrations are as described in Fig. 7

The difference of responses in the presence and absence of Ng are explained by its combined effects on CaN and CaMKII, different according to the frequency (Fig. 9b). The absence of Ng increases both CaMKII and CaN activation at low spike frequencies, however the effect on CaN is far greater, thus favouring LTD. As the spike frequency increases, the activity of both enzyme increases. However, while CaMKII continues to increase until reaching a plateau, CaN activity peaks and then decreases, shifting the effect from LTD to LTP. The peak occurs at lower frequencies in the absence of Ng (3 Hz instead of 10 Hz), explaining the left-shift of BCM curve. At high frequencies, the effect of Ng is opposite, in direction and intensity, on both enzymes. The CaMKII plateau is increased three-fold in presence of Ng. On the contrary, CaN plateau is only slightly decreased by Ng. The net effect is a much stronger overall kinase activity in presence of Ng.

Therefore, Ng regulates the activation of CaMKII and CaN, thus modulating synaptic plasticity in three phases: At low spike frequency, Ng inhibits the activation of both proteins, however exerts a stronger effect on CaN, thus inhibiting LTD. At intermediate frequencies, Ng inhibits both enzyme, albeit exerting a stronger effect on CaMKII, thus delaying the onset of LTP. At high frequencies, Ng inhibits CaN but facilitates CaMKII autophosphorylation thus facilitating LTP. These phases might provide explanations for opposing experimental observations on LTP induction in Ng knock-out mouse. Neurogranin raises the threshold frequency to activate CaMKII, however, once the threshold is crossed, neurogranin facilitates its activation.

### Neurogranin facilitates CaMKII activation via the inhibition of CaN

Neurogranin’s positive impact on CaMKII activation at high frequency electric stimulations has been observed experimentally [15, 19], but no underlying mechanisms were suggested. As in the absence of Ng, decreased CaMKII activation is composed of phosphorylated CaMKII monomers, and CaN is responsible for dephosphorylating CaMKII, we hypothesized that CaN might play an important role in mediating Ng’s effect on synaptic plasticity.

The frequency response plot (Fig. 9b) shows that the absence of Ng causes a large change of the CaMKII plateau at high frequencies, but a fairly small change of CaN’s. It is therefore difficult to see how such a small change of CaN activity could have an important effect on CaMKII autophosphorylation. One thing to bear in mind is that activated-areas, calculated at each frequency, represent the activity above a basal level, as it characterizes the amplitude and duration of the response to the calcium stimulation. However, the inhibition on the CaMKII autophosphorylation is not only due to the dynamics of CaN following calcium signals, but to CaN basal level. The latter is elevated in the Ng-absent situation (Fig. 7a,b). Taking into consideration this increase would cause a vertical shift of about 30 units in its frequency dependence curve (Fig. 9b green crosses) without affecting the CaMKII curve much. Ng presence therefore changes the situation dramatically. CaN activity is higher than CaMKII activity in the absence of Ng, whereas the opposite situation is true when Ng is present.

To further assess CaN’s role, we removed it from the model and re-simulated the various frequencies of calcium spikes, with or without Ng. As shown in Fig. 10a, removing CaN dramatically increases CaMKII activity at high calcium spike frequencies regardless of the Ng’s presence. Moreover, CaMKII activities reach the same maximums, Ng’s positive impact being completely lost. The major change upon removing Ng is the shift towards lower spike frequencies shows easier onset of LTP at intermediate frequencies. This indicates that, without CaN, the only impact of Ng on CaMKII activation is negative.

**Fig 10.**
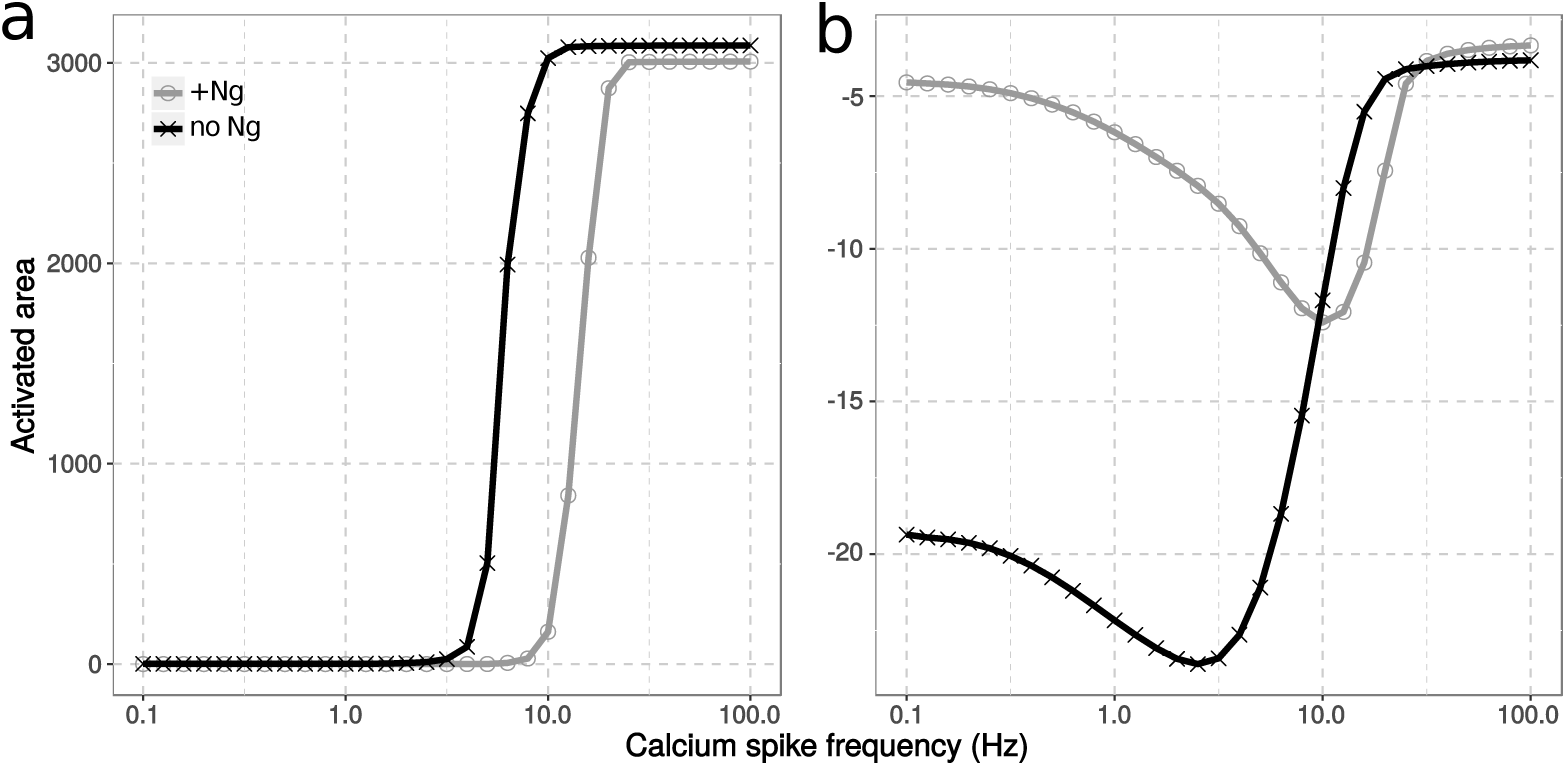
Calcineurin mediates neurogranin’s effect on CaMKII activation. Models were simulated without CaN (a) or with double amount of CaN (b). The combined response of CaMKII and CaN to calcium spike frequencies were quantified as described in Fig. 9, with (grey circle) or without neurogranin (black cross). Simulation conditions and concentrations are as described in Fig. 7. Double amount of CaN equals to 16 µM.

We then doubled CaN concentration, to twice its detected level in hippocampus CA1 region [59] (Fig. 10b). At low spike frequencies, CaN activity wins over CaMKII’s as seen above with normal CaN concentration. However, at high frequencies, CaMKII activity can no longer overturn CaN activation resulting in abolished LTP induction regardless of Ng’s presence. This indicates that with high CaN concentration, Ng produces no effect on CaMKII activation nor facilitating LTP induction. To summarize, when there is not enough or too much CaN, CaMKII activation is either excessive or deficient. In either cases, Ng exerts no positive effect on CaMKII activation, other than limiting calmodulin’s availability.

We tested other CaN concentrations and plotted the differences of CaMKII activation reached at the high calcium spike frequencie (100 Hz), in presence and absence of Ng. Fig. 11 shows a non-linear relationship between the concentration of CaN and the effect of Ng on CaMKII activation. It seems that with our concentrations of Ng, CaM and CaMKII, Ng displays the highest positive effect on CaMKII activation when CaN concentration is about 4 micromolar. Either reducing or increasing CaN concentration weakens Ng impact.

**Fig 11.**
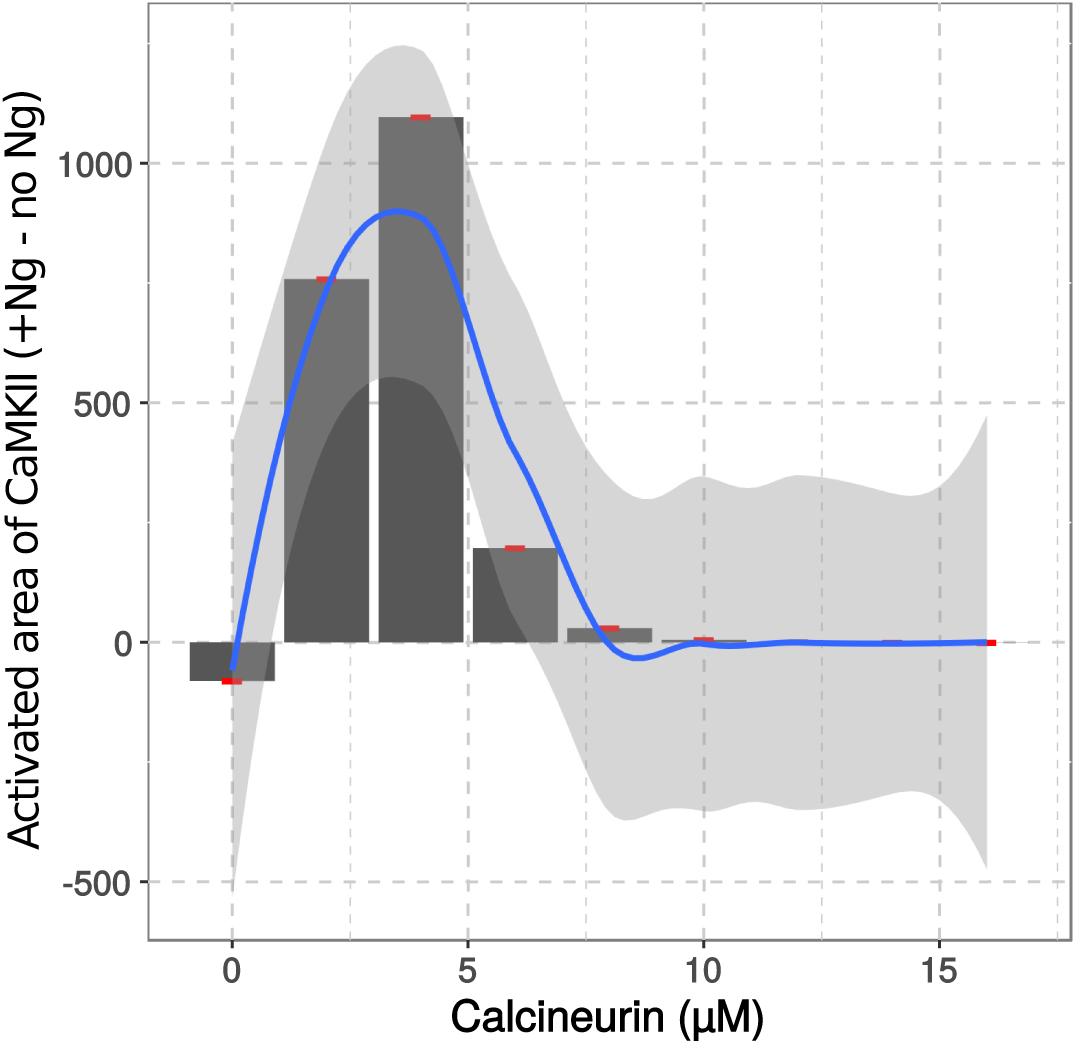
Calcineurin concentration affects neurogranin regulation of CaMKII activation. Calcineurin concentration was varied in presence (wt) and absence (NgKO) of Neurogranin. The differences of CaMKII’s activated area reached in response to 100Hz calcium spike frequency between wt and NgKO situation were plotted against CaN concentration. The relationship was further fitted into a polynomial function (method loess() from ggplot2) and was plotted as the blue line. Red points are the values from our simulations. Shaded area labels the 95% confidence interval for the fitting.

These results suggest a central role for CaN in controlling how Ng affects CaMKII activation at higher calcium spike frequencies. The presence of neurogranin facilitates a prompt release of large quantities of calmodulin at specific spike frequencies, which facilitates the onset of CaMKII activation and autophosphorylation, allowing it to overcome CaN’s negative feedback. Removal or over-expression of CaN remove neurogranin’s positive regulation on CaMKII, reducing its role to merely a calmodulin buffer.

### The concentration of neurogranin impacts LTP induction

Previous research showed that Ng undergoes constant shuffling in and out of the PSD. Its function as a regulator of synaptic plasticity may rely on its anchoring in the PSD and the resulting recruitment of more calmodulin [60]. While this is certainly an important aspect of Ng function, modelling Ng and calmodulin translocation in and out the PSD may blur other Ng impacts on plasticity. We have already shown that calmodulin concentration is a limiting factor in the PSD and increasing it has positive effect on LTP induction [10]. Although our current model only incorporates neurogranin binding of calmodulin in an homogeneous compartment, we can already show that increasing Ng concentration, while keeping calmodulin concentration intact, has a positive effect on LTP induction. As shown in Fig. 12, increasing Ng concentration results in a larger activation of CaMKII over CaN, with a steepest transition from CaN to CaMKII as we increase calcium spike frequency, and higher plateaus for CaMKII activity at both low and high frequencies. Despite this relative shift toward CaMKII at all frequencies, the transition between CaN and CaMKII (Θ_*m*_) occurs at higher frequencies when Ng concentration increases.

**Fig 12.**
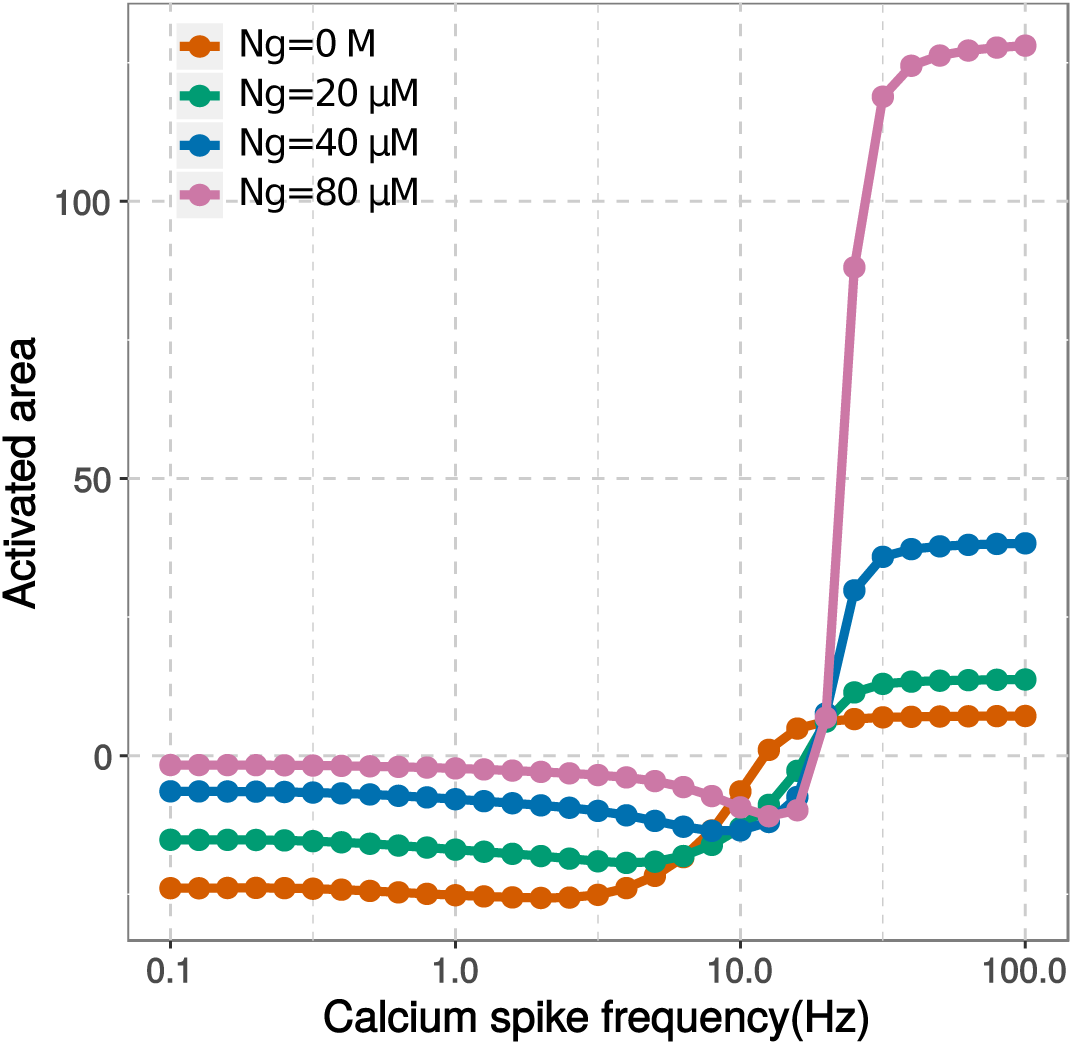
Neurogranin’s concentration affects synaptic plasticity. Computational models with various neurogranin concentration were stimulated by 300 calcium spikes at various frequencies. The activated area of CaMKII and CaN were calculated and their net effects on AMPA receptor phosphorylation were calculated as described in Fig.7, and plotted as a function of calcium spike frequency.

The level of Ng in the brain, especially in the hippocampus, is declining with age [61–63]. Our simulation results indicate a direct relation between the concentration of Ng and the activity level of CaMKII, which may provide explanations for the deteriorating cognitive functions among the ageing population.

## Discussion

Calmodulin’s impact on synaptic plasticity are mediated via and regulated by its binding partners [26, 31, 64]. Contrarily to enzymes which downstream activities are well documented, the role of some of the non-catalytic proteins is less clear. One such protein is neurogranin, a highly expressed brain protein that carries a calmodulin-binding IQ domain [65, 66].

Neurogranin primarily interacts with the closed conformation of calmodulin C-lobe [13, 26], and is considered to buffer calmodulin [11, 12, 66, 67]. Despite many studies of its involvement in regulating synaptic plasticity, its precise role is still questioned. Several *in vivo* experiments showed that neurogranin facilitates CaMKII activation during LTP induction protocols [16, 19], and is not merely a calmodulin buffer, as CaMKII activation requires a significant amount of calmodulin in the open conformation. To confuse matters further, neurogranin knock-out in mouse can either facilitate or inhibit LTP induction [14–19].

By setting up a mechanistic model of calmodulin conformational changes of its two lobes, we explored the potential roles neurogranin plays in regulating bidirectional synaptic plasticity mediated by CaMKII and calcineurin.

We firstly examined how Ng coordinates calmodulin lobes in response to calcium spikes frequencies. It has been established that calmodulin C lobe, although having higher affinity, has slower binding kinetics for calcium ions than the N lobe [11, 20, 23, 31]. Therefore, it has been proposed that the calcium binding to C lobe is rate limiting for calmodulin opening [26, 32, 64]. However, our simulation results showed the contrary. When no calmodulin binding partner is presented, calmodulin N lobe is harder to open than C lobe, requiring higher calcium spike frequencies and larger free calcium elevations (Fig. 6a). This is because the fast binding and dissociation kinetics of N lobe make it follows calcium spikes too close to open before releasing calcium. We futher elucidated that the role Ng plays is to reduce these differences between calmodulin lobes and to synchronize their opening at specific calcium spike frequencies. This is achieved by Ng’s preferential binding to the C lobe and limiting its opening at low frequencies (Fig. 6b). The advantage of this synchronization is to minimize calmodulin opening when calcium stimulation is weak, therefore reducing basal calcineurin activity and shifting the majority of opened calmodulin to CaMKII when calcium signal is strong (Fig. 7, 9).

Our findings are in agreement with some studies that have proposed that Ng and other IQ motif proteins act as calmodulin caches, enhancing the activation of calmodulin binding partners when calcium concentration is high [41, 65, 68]. Our study went a step further by showing that this increased calcium concentration is the consequence of increased calcium spike frequencies, other than total calcium ion inputs. Furthermore, the mechanisms underlying this dynamic regulation is rooted in the allosteric regulation of calmodulin lobes and the reciprocal influence among calmodulin and its binding partners.

Our simplified frequency response curve, that was drawn from the differences of integrated enzymatic efficiencies between CaMKII and CaN (Fig. 9a), matches with the experimental observations by Huang *et al.* [16]. The absence of neurogranin facilitates LTD but impairs LTP induction, resulting in a down-shift BCM curve. In addition, we observed a slight left shift when Ng was removed, indicating a potential easier onset of LTP at these intermediate frequencies, dependent on the threshold CaMKII required. The major experimental study questioning Ng’s positive role on synaptic plasticity is published by Krucker *et al.* [19]. In their study, the mouse brains that are absent from functional Ng display lowered CaMKII activity, but are enhanced to exhibit LTP under high frequency stimulations [19]. Our research might shed light on this discrepancy, as we showed that the absence of Ng does not abolish CaMKII activation at high intensity calcium stimulation and most importantly, enhances CaMKII activation at intermediate frequencies. With the optimal amount of CaN, CaMKII activity can overturn CaN activity at lower frequency in the absence of Ng than when it is present, therefore potentially facilitating LTP induction. We further elucidated that the LTP induction frequencies are likely to be dynamically regulated by protein expression level, largely due to neurogranin itself (Fig. 12).

Perhaps, the most intriguing finding from our research is that CaN facilitates Ng to enhance LTP induction. CaN is one of the highly expressed protein phosphatases in nervous system [69], and its dysfunction has been associated with many neurological diseases [70–73]. CaN has been shown to be involved in processes weakening synaptic connections [74–79]. And blockage of CaN has shown to encourage learning and memory [80, 81].

Our finding do not contradict with these views. In fact, we showed that neurogranin positive impact on CaMKII activation and LTP induction is due to its inhibition of CaN, especially at the basal calcium concentrations. When Ng is knocked out, calmodulin is prone to be activated by basal levels of calcium, therefore elevating the resting level of CaN activity, which in turn decreases CaMKII autophosphorylation. Therefore, when CaN concentration is too low, the difference of CaN basal activity in presence or absence of Ng has very little impact on CaMKII activity. On the other hand, when CaN concentration is high, even in presence of Ng, CaN can still binds to most CaM, preventing CaMKII-CaM binding and subsequent CaMKII autophosphorylation.

This sensitivity to CaN concentration may provide further explanation for conflicting experimental observations where Ng knock out does not always result in reduced LTP induction [15, 19]. Further experimental validation focusing on CaN’s contribution to Ng role on synaptic plasticity would definitely provide new insights about Ng function and better understanding of synaptic plasticity.

In both bursting and continuous stimulations, high frequency calcium stimulations only produce a small amount long lasting active CaMKII monomers. Because of the large total number of monomers included in the model, this still corresponds to approximately 700 monomers, or nearly 60 CaMKII dodecamers. Since NMDA receptors, especially the NR2B-containing ones, are present in the PSD in small quantities [82], these active CaMKII dodecamers can be sufficient to saturate the C terminal tail of NR2B containing NMDA receptors, therefore facilitating AMPA receptor insertion and LTP induction.

This also reveals a limitation in our current approach to approximate CaMKII autophosphorylation rates, that lowers the probability for a given monomer to have an active neighbour. Our dendritic spine contains a very large amount of CaMKII monomers, representing a large number of hexamers. Since we assume a single homogeneous compartment, all monomers possess an equal chance to interact with calmodulin. This increases the amount of active calmodulin required to obtain two adjacent CaMKII monomers in a given hexamer ring, a requirement for trans autophosphorylation. An improved approach would be to consider the PSD as a spatially confined compartment segregated from the rest of the spine, and to incorporate CaMKII dodecamer clustering and transport in and out of this compartment. This would increase the chance of PSD-located CaMKII to bind calmodulin, and would potentially provide a more accurate estimation of autophosphorylation rates. However, the current low estimation does not affect our main conclusion, which compares CaMKII activation in presence and absence of Ng since CaMKII autophosphorylation rates are adjusted the same way in both situations. The only difference could be that the frequencies we found to be able to induce LTP might be higher than reality, albeit for both scenarios.

We have taken a highly simplified concept to build our model upon, in which the effectiveness of the phosphatase and the kinase is directly used as a measure for synaptic plasticity. Although the requirements for these enzymes and their central roles in synaptic plasticity are well established [9, 38], there are more modulators and interactions in these processes and at different stages of memory consolidations [9]. The difficulties of incorporating more molecular interactions in the model are estimating their relevant equilibrium and kinetic parameters systematically, and ensuring their identifiabilities, requirements we tried to ensure in this study. Our mathematical model and parameters can be reused and expanded to further our understanding of calmodulin and its binding proteins.

In conclusion, our study revealed a complex and dynamic role neurogranin plays in regulating bidirectional synaptic plasticity. It functions via preferential binding to calmodulin C lobe at the closed comformation. And by doing so, Ng synchronizes the two calmodulin lobes opening, limits CaMKII activation at intermediate calcium spike frequencies, while facilitating it at high frequencies. Apparently contradictory experimental observations regarding its effect on LTP induction might be all valid. As we already showed before [10], the exact CaMKII activity required to induce LTP is dependent on conditions and other molecular components inside the post-synaptic dendritic spine. Ng’s positive regulation of CaMKII activity depends on both its concentration and that of CaN. Furthermore, it depends on calcium spike frequencies and the threshold CaMKII activity required to facilitate AMPA receptor insertion and LTP induction.

## Supporting Information

**S1 Fig. Estimations of boundaries for allosteric parameter cC and calcium affinity to a binding site on calmodulin’s C lobe (KCT) and N lobe (KAT)**, **as well as affinity between CaMKII peptide and open calmodulin (Kd CaMKIIpeptide RR).**

**S2 Fig. Estimations and correlations of allosteric parameters cC**, **cN**, **LC**, **LN**, **and affinity between neurogranin and closed calmodulin (Kd Ng TT).**

**S3 Fig. Steady state validation of calcium binding calmodulin.** Comparison of steady state simulation results (solid lines) and published experimental observations (dots) of calcium binding C lobe (a) and N lobe (b) of calmodulin. To compare with experimental data published by Hoffman *et al.* [26], the model was set up with [CaM] = 5 µM and [Ng] = 50 µM. To compare with experiments published by Evans *et al.* [6], model initial states were set up as [CaM] = 2 µM and [CaMKII] = 10 µM.

**S4 Fig. Linear relationship between log10(cC) and log10(LC).**

**S5 Fig. Estimations of calcineurin binding (kon PP2B CaM) and releasing open calmodulin (koff PP2B CaM).**

**S6 Fig. Estimation of allosteric parameter cC and association constants for calmodulin binding CaMKII (kon CaMKII)**, **phospho-CaMKII (kon CaMKIIp) and neurogranin (kon Ng)**, **calcium binding C lobe of calmodulin (kon CT) and the conformational transition rate of C lobe**, **when no calcium or protein is bound (k T2R C0).**

**S7 Fig. Correlation analysis of each pair of the parameters estimated above in S6 Fig**, **based on parameter values searched and their scores for fitting experimental observations.**

**S8 Fig. Validations of calcium dissociation kinetics from calmodulin.** Calcium dissociation from C-lobe of calmodulin (a,b) were measured upon mixing calcium chelator EGTA (10^−2^ M), and simulated after reaching equilibrium with pre-mixed calmodulin (10 µM) and calcium (100 µM) with (cyan) or without (salmon) neurogranin (50 µM). Calcium dissociation from whole calmodulin (c) were measured upon mixing calcium chelator quin2 (150 µM), and simulated after reaching equilibrium with pre-mixed calmodulin (2 µM) and calcium (20 µM) with CaMKII (2 µM salmon) or phospho-CaMKII (2 µM cyan). Solid line: simulation results; dots: experimental observations [26, 31].

**S9 Fig. Calculation of the rate of CaMKII autophosphorylation.** The procedure for calculating the rate of CaMKII autophosphorylation rate as a function of active CaMKII monomers (for details see Methods section).

**S10 Fig. Protein activations after calcium inputs at 30 Hz.** Computational models with (a) or without (b) neurogranin were stimulated by 300 calcium spikes at 30Hz. Protein activities were normalized by their total concentration, and CaMKII and CaN were further multiplied by their catalytic efficiencies towards GluR1 subunit of AMPAR. [CaM] = 40 µM, [Ng] = 40 µM, [CaN] = 8 µM, [CaMKII] = 80 µM; *kcat*_CaMKII_ = 2 s^-1^; *kcat*_CaN_ = 0.5^-1^

**S11 Fig. Protein activations after calcium inputs at 100 Hz.** Computational models were simulated with (a) or without (b) neurogranin, with 300 calcium spikes at 100 Hz. Each calcium input was as described in Fig.3. Both protein activities were normalized to their total concentration, then multiplied by their catalytic constant for GluR1. [CaM] = 40 µM, [Ng] = 40 µM, [CaN] = 8 µM, [CaMKII] = 80 µM; *kcat*_CaMKII_ = 2 s^-1^; *kcat*_CaN_ = 0.5^-1^

## Acknowledgments

We thank Dr Len Stephens, Dr Simon Andrews, and Dr Michael Wakelam for their support and help.

